# Acquisition and reversal of glioblastoma chemoresistance are mediated by the Rho GTPase pathway

**DOI:** 10.1101/2024.10.23.619654

**Authors:** Yuli Thamires Magalhaes, Viktor Kalbermatter Boell, Fabio Luis Forti

## Abstract

Glioblastoma (GBM) are highly aggressive tumors treated mainly with surgery, radiotherapy, and chemotherapy. Innovative multimodal therapies are needed, targeting the immune system, tumor metabolism, and cell signaling. Our research focuses on the role of the actin cytoskeleton and Rho GTPases in modulating DNA damage repair and therapeutic sensitivity in GBM cells. We developed GBM sublines resistant to temozolomide (TMZ) and cisplatin (CP), and assessed actin stress fiber organization, Rho pathway activity, and resistance phenotype. TMZ-resistant clones exhibited increased Rho pathway activity, elevated p53 and DNA double-strand break (DSB) repair pathways, but reduced MMR protein levels. Importantly, Rho GTPase inhibition restored TMZ-resistant clones’ sensitivity to TMZ and CP, counteracting chemoresistance. While both drugs reduced DNA repair capacity in normal GBM cells—exacerbated by Rho inhibition—TMZ-resistant clones with overactivated Rho pathways did not show this effect. This response was p53-wild-type dependent, as p53-mutant GBM cells were unresponsive to Rho inhibition. However, p53-mutant cells treated with PRIMA-1 showed restored sensitivity to chemotherapeutics with Rho inhibition. Furthermore, modulation of the actin cytoskeleton and Rho GTPases affected sensitivity and viability in GBM spheroid models exposed to chemotherapy. In summary, Rho pathway activity and actin cytoskeleton dynamics are critical for both the development and reversal of chemoresistance in GBM tumors.

**STATEMENT OF SIGNIFICANCE:** Chemoresistance in glioblastomas modulates the Rho GTPases pathway and actin cytoskeleton, while negatively affecting DNA repair. Downmodulating the actin circuitry in resistant GBMs sensitizes them to TMZ and CP drugs.

## INTRODUCTION

Rho GTPases, a family of small signaling G proteins comprising Rho, Rac, and Cdc42 among others, are critical regulators of the actin cytoskeleton dynamic, acting in a variety of cellular process associated with carcinogenesis, including cell migration, invasion, division and metastasis (1). Actin cytoskeleton remodeling is a critical event in cancer progression, enabling tumor cells to adopt more invasive phenotypes and resist apoptotic signals. Rho GTPases act as molecular switches that integrate signals from the extracellular matrix and intracellular pathways, orchestrating changes in the actin network that contribute to cancer cell survival and chemoresistance (2). Furthermore, Rho GTPase-mediated regulation of the actin cytoskeleton can impact the effectiveness of chemotherapeutic agents by altering drug uptake, promoting survival pathways, and fostering the formation of drug-resistant cell populations (3).

In glioblastoma multiforme (GBM), the most incident and malignant brain tumor, with a grim prognosis mostly due its ability to develop resistance to chemotherapeutic agents, Rho GTPases have emerged as key players in maintaining the tumor’s malignant phenotype. Rho/actin signaling has been implicated to the enhanced tumor’s invasive capabilities and the ability to resist apoptosis induced by chemotherapeutic agents, which contributes to its resistance (4). These treatment-resistant phenotype is largely facilitated by robust DNA damage response (DDR) mechanisms and DNA repair pathways, processes where Rho GTPases has been recently related to, promoting the recognition and repair of DNA double-strand breaks (DSBs) (5,6). Particularly, the interplay between Rho GTPases and the tumor suppressor p53 has attracted significant interest. Wild-type p53 is pivotal in regulating cell proliferation, survival, and apoptosis, but mutations in p53—observed in 40% to 50% of GBM cases—disrupt these functions, leading to enhanced tumor resilience (7). These mutations often result in a loss of p53’s transcriptional activity, contributing to the tumor’s resistance to DNA-damaging treatments (8).

Recently, our group elucidated the mechanism by which downregulation of the Rho GTPase pathway abrogates resistance to ionizing radiation in wild type p53 glioblastoma by suppressing DNA repair mechanisms (9). We demonstrated that inhibiting the Rho GTPase pathway increases the sensitivity of GBM cells to ionizing radiation by increasing the number of DNA double-strand breaks and delaying their repair through non-homologous end-joining (NHEJ). Additionally, we found that the Rho GTPase pathway intercommunicates with the p53 pathway through nuclear translocation of p53 facilitated by G-actin, influencing DDR signaling. This interdependence suggests that targeting the Rho GTPase pathway could effectively sensitize GBM cells to radiotherapy and potentially overcome acquired resistance (9).

This paper aims to explore whether the molecular underpinnings of Rho GTPase-mediated radioresistance in GBM are also mediating the chemoresistance of these tumors, and to present evidence for the involvement of modulation of DNA repair mechanisms in this therapeutic response. To this end, we generated temozolomide (TMZ)- and cisplatin-resistant p53 wild-type GBM sublines and examined potential changes in the Rho and DDR pathways during the acquisition of resistance. By targeting the Rho GTPase pathway in these resistant GBM cells, as well as in relative spheroid tumor models, we may propose new avenues to increase the efficacy of existing GBM treatments and overcome chemoresistance concomitantly with radioresistance.

## MATERIALS AND METHODS

### Glioblastoma cell lines culture condition

Human glioblastoma cell lines, including p53 wild-type (U87-MG, A172, U343-MG) and p53 mutant (T98G, U138-MG, U251-MG), were sourced from the American Type Culture Collection (ATCC) and provided by Prof. Dr. Menck (ICB-USP), Prof. Dr. Sogayar (IQUSP and FMUSP), and Prof. Dr. Hojo (FFCLRP-USP). Cells were cultured as previously (9) in DMEM supplemented with 10% FBS, 25 μg/mL ampicillin, and 100 μg/mL streptomycin at 37°C in 5% CO_2_. Subculturing occurred every 48-72 hours for up to 4 weeks, with regular monitoring for genotypic, phenotypic, and mycoplasma contamination to ensure the quality of the model.

### Drugs and treatments

Chemotherapeutic agents included cisplatin (CP, Sigma-Aldrich) and temozolomide (TMZ, Sigma-Aldrich). Stock solutions of CP (10 mM in 0.9% NaCl) and TMZ (50 mM in DMSO) were stored at 4°C and -20°C, respectively. Prior to use, CP was warmed to 60°C. Rho pathway inhibition by C3 toxin was achieved through transfection (Lipofectamine 3000, Invitrogen) of pEF-myc-C3. F-actin disruption was induced by Cytochalasin D (CD, 100 nM, Sigma–Aldrich). Knockdown of wild-type p53 was performed using specific siRNA (MISSION® esiRNA, Invitrogen), and p53 mutant activity was restored using PRIMA-1 (PR-1, 25 µM, Sigma–Aldrich) for 24 hours (9).

### Spheroid formation from GBM cells

Spheroids were prepared using the liquid overlay technique (10). In brief, U87-MG cells (1 to 3 x 10^4^ cells) were seeded on agarose-coated 96-well plates (50 µL of 1.5% agarose/well) and cultured as described. Spheroid formation and integrity were evaluated after three days. Morphometric parameters were acquired using a phase-contrast microscope and analyzed with ImageJ. Treatments involved replacing half of the media with fresh media containing double the desired drug concentrations. Spheroid viability was assessed using the acid phosphatase (APH) assay (10).

### Rounds of CP and TMZ exposure for increasing drug tolerance

To study the effects of prolonged chemotherapeutic exposure, U87-MG cells were treated with increasing concentrations of CP (0.25 µM to 1.5 µM) or TMZ (2.5 µM to 20 µM) over two rounds. Each round involved a 72-hour drug exposure followed by a 7-day recovery for CP or 4-day recovery for TMZ. Chemoresistance development was monitored via cell viability, phalloidin staining, and levels of Rho, p53, and DNA repair proteins, assessed by immunofluorescence and immunoblotting.

### Generation of chemoresistant resistant clones

Chemoresistant clones were generated using a rigorously established protocol, by treating cells with increasing doses of TMZ as detailed above. Cells were divided into subgroups (A to I), each one subjected to specific TMZ doses for 72 hours followed by a 4-day recovery. This cycle was repeated with escalating TMZ concentrations until resistance development. Resistant clones were expanded, cryopreserved, and validated by subsequent viability assays.

### Cell viability assays

For IC_50_ determination and chemoresistance evaluation in monolayers, cell viability was assessed using the MTT assay (9). Briefly, cells were treated for up to 72 hours, and MTT reagent (0.5 mg/mL, Invitrogen) was added, followed by a 3-hour incubation. Formazan crystals were dissolved in DMSO, and absorbance was measured at 570 nm using a plate reader (Epoch, BioTek). For spheroids, IC_50_ was determined using a modified APH assay (9,10), in which a 96-well conical-bottom microplate was used to replace the centrifugation steps, with absorbance measured at 405 nm. IC_50_ values (µM), determination coefficient (R²), and 95% confidence intervals (CI) were determined using nonlinear regression of dose-response curves in GraphPad Prism 10.

### Cell proliferation assay

Cell proliferation was monitored by seeding 1 x 10^4^ cells in 12-well plates, followed by daily cell counts for 8 days using a Neubauer chamber. C3 transfection occurred 24 hours post-seeding, with subsequent TMZ or CP treatment 6 hours post-transfection. Doubling time and 95% CI were determined using nonlinear regression of growth curves in GraphPad Prism 10.

### Cell survival assay

Colonies were obtained by seeding 1 x 10^3^ cells. in 6-well plates, 24 hours prior to treatment exposure. After 10-14 days, colonies were fixed with 10% formaldehyde, stained with 5% crystal violet, and counted to determine survival fractions.

### Rho activity assay and immunoblotting

Rho activity was measured using a pulldown assay as previously described (9). Briefly, cellular protein extracts were incubated with glutathione-Sepharose beads bound to the RBD-GST fusion protein. Immunoblotting was performed on 25 µg of protein lysate, quantified with Bradford reagent (BioRad), denatured with the Laemmli protocol, separated by SDS_JPAGE, and transferred to nitrocellulose membranes (Millipore). Membranes were blocked in 5% low-fat milk for 30 min, incubated with specific primary and secondary antibodies (Table S1) , and quantified using ImageJ.

### Immunofluorescence assay

For immunofluorescence (9), cells were seeded on glass coverslips, fixed with 4% paraformaldehyde, permeabilized, and stained with Phalloidin and/or primary antibodies, followed by Alexa Fluor-conjugated secondary antibodies (Table S2). Fluorescence was visualized using a confocal microscope (Leica Microsystems), and nuclear p53 was quantified using the TissueFAXS i-Fluo system (TissueGnostics) and StrataQuest software.

### Host Cell Reactivation (HCR) assay for CP- and TMZ-Induced DNA damage

HCR assays were conducted as previously described (11,12) with modifications. Briefly, pLuc (firefly luciferase) plasmids were treated with 750 nM CP or 100 nM TMZ for 4 hours at 37°C before being transfected into cells seeded in 96-well white plates. Co-transfection of pLuc and pRenilla (transfection control), with or without pEF-myc-C3, was performed using Lipofectamine 3000 reagent according to the manufacturer’s instructions. Luciferase activity was measured 48 hours post-transfection using the Dual-Glo Luciferase Assay System (Promega). DNA repair capacity was expressed as a percentage of luciferase activity relative to untreated plasmids.

### Statistical Analysis

Experiments were conducted with at least three biological replicates, each with two to six technical replicates. Experiments Statistical significance was assessed using two-way ANOVA with post hoc tests (Tukey’s or Sidak’s) or one-way ANOVA for two-group comparisons, using GraphPad Prism 10. Statistical significance was denoted as ∗p ≤ 0.05, ∗∗p ≤ 0.01, ∗∗∗ p ≤ 0.001, and ****p < 0.0001.

## RESULTS

### Involvement of Rho pathway in the chemotherapy response of glioblastomas

Chemotherapy drugs such as TMZ and CP causes, respectively, on DNA: induction of O^6^-methylguanine (O^6^-MeG), which ultimately leads to the formation of DNA double-strand breaks (DSB), and purine adducts, which can lead to intra and interstrand crosslinks (ICLs). This study on the regulatory mechanism of the Rho pathway in the DNA damage repair caused by chemotherapy drugs started by interrogating gene expression databases, on the EMBL Cell Expression Atlas platform (https://www.ebi.ac.uk/gxa/home). Thus, a bioinformatic analyses of gene expression was performed for the wild type p53 U87-MG glioblastoma cell line treated with low doses of CP (E-GEOD-66493 study) at the timepoints of 6 and 24 hours (Fig. S1A); raw data were obtained in Log_2_(fold change) between the treated conditions and the basal gene expression. The short treatment with CP led to a smaller response in relation to gene expression, presenting only 42 differentially expressed genes while the prolonged treatment led to the differential expression of 380 genes, indicating a greater cell response to the accumulation of CP adducts (Fig. S1B). Through a set analysis it is possible to see that most of the genes initially affected by the short treatment with CP remain with their altered expression during the prolonged treatment (Fig. S1C). Experimental validation conducted with treatments with CP or TMZ, at the same time points, showed that they similarly triggered DNA damage response causing p53-Ser15 phosphorylation with subsequent p21^Kip1^ expression in U87-MG cells. In agreement with datamining, Rho GTPase pathway is also activated at these conditions as seen by cofilin-1 (CFL1) phosphorylation at Ser3, especially in the prolonged exposure to chemotherapeutics (Fig. 1A). Trying to establish a tentative protocol for generating chemoresistant U87-MG cells after two rounds of prolonged treatments with different and low doses of CP or TMZ, two sets of samples after the respective recovery time points were collected (Fig. 1B). It is noted that chemoresistance start to show up already after the second round of treatment, where cells displayed greater viability in response to higher doses of CP or TMZ (Fig. 1C). The samples extracted at the end of each treatment round were submitted to immunofluorescence assays to observe the actin cytoskeleton integrity and cell morphology. Prolonged exposure to different doses of CP (Fig. 1D) or TMZ (Fig. 1E) altered the structure of the actin filaments, leading to the formation of stress fibers, protrusions, lamellipodia and filopodia (white arrows). For both drugs, the first treatment round led to greater formation of stress fibers and some points of F-actin clusters, in addition to the formation of long and thin protrusions, being these effects more evident in treatments with higher doses. The second treatment round also altered the cellular morphology, with a greater frequency of formation of protrusions and stress fibers. Immunoblotting data from the samples extracted after the two rounds of CP treatments (Fig. S2), showed that first exposure to CP activates the p53 pathway, evidenced by its phosphorylation and p21^Kip1^ expression, as well as Rho signaling activation shown by cofilin 1 phosphorylation. Most interestingly was that the second round showed a strong abrogation of the p53 pathway while the Rho pathway persisted active. Activation of Rho pathway and cytoskeleton remodeling, visualized in the morphological analysis, caused by CP (or TMZ) agree with the genes differentially expressed after 6h and 24h of CP exposure. For that, pathway enrichment was analyzed by the EnrichR platform and among the main pathways affected in response to CP there was the p53 (and its homologs p63 and p73), cell cycle regulation and DNA damage response pathways (Fig. S1D) commonly activated in response to genotoxic stress. Short exposure to CP (Fig. S1E) led to modulation of similar response pathways, while long-term exposure to CP showed more interesting outcomes that included the Rho GTPase pathway (Fig. S1F). These results show that short-term treatment modulates p53 signaling pathways, while long-term treatment modulates Rho GTPase pathways in addition to p53 pathways, indicating an intricate relationship between the two pathways in response to chemotherapy drugs.

**Figure 1.**
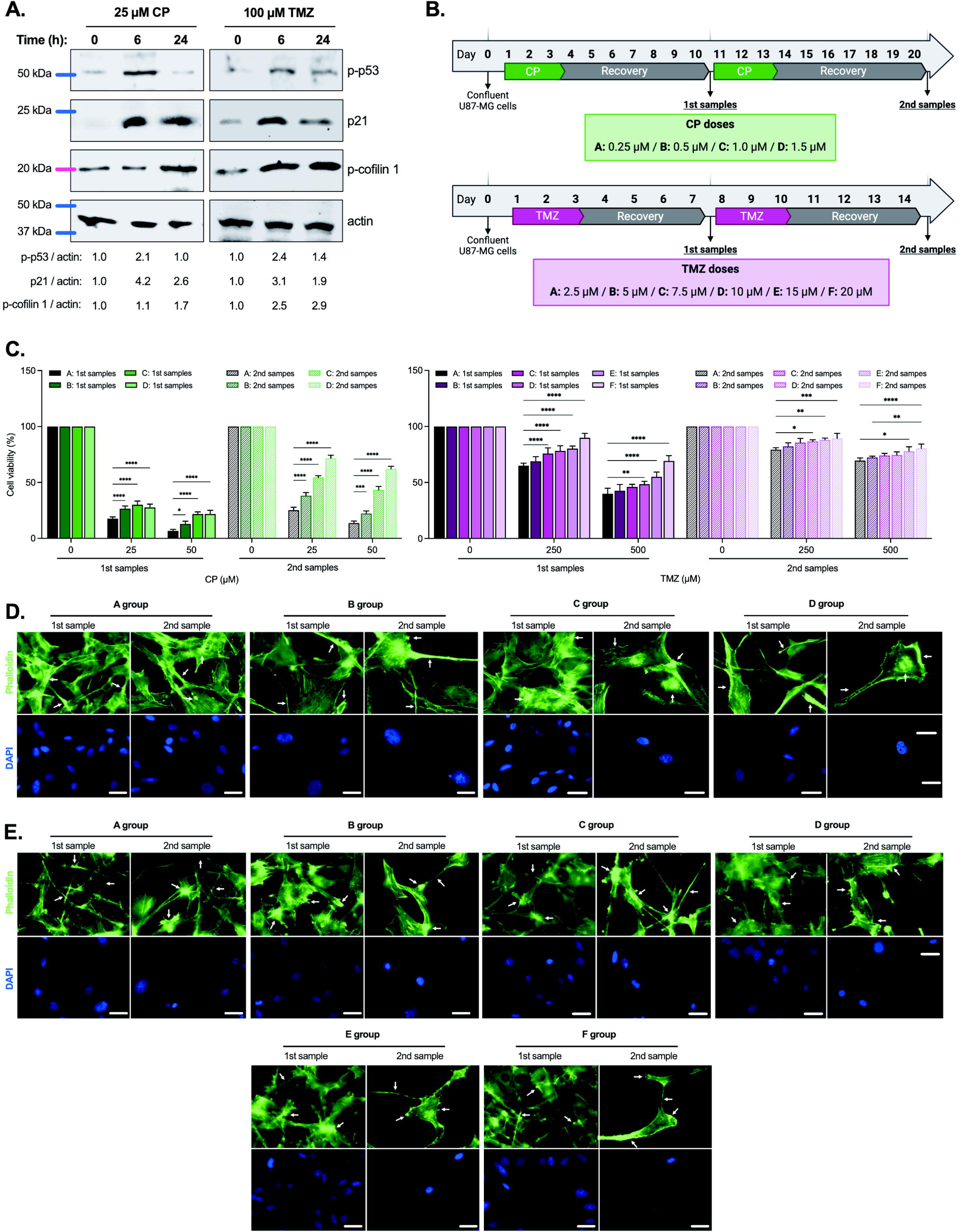
The prolonged exposure of GBM cells to chemotherapeutics is accompanied by alterations in the Rho pathway and F-actin structures. **A.** Representative image of immunoblotting assays from U87-MG cells exposed to CP 25 µM or TMZ 100 µM for 6h and 24h, showing the levels of p21, p-p53, and p-cofilin 1. **B.** Schematic representation of treatment protocols involving two rounds of treatment/recovery with varying doses of CP or TMZ in U87-MG cells for the specified duration. For each drug, cells were divided into groups (A–D for CP, A–F for TMZ) according to the indicated drug concentrations. Cells were harvested after the first (1st samples) and second rounds of treatment (2nd samples) and subjected to viability and phalloidin staining assays. **C.** MTT viability assays of U87-MG cell groups following CP (green) or TMZ (pink) treatment protocols and collected at different timepoints. Cell groups were treated with CP (25 or 50 µM) or TMZ (250 or 500 µM) for 72 h. **D-E.** F-actin staining using Alexa Fluor 488-phalloidin of U87-MG cell groups following CP (**D**) or TMZ (**E**). The images were captured with a 63x magnification objective and a scale bar of 25 μm.

### Generation of TMZ-resistant U87-MG cell clones

To obtain chemoresistant cell clones in vitro, mimicking clinical resistant or recurrent tumors, we developed a treatment regimen strategy that worked perfectly for TMZ (Fig. 2A), but unfortunately not for CP. Thus, using different rounds of treatment/recovery for reaching a different final cumulative dose of TMZ (Fig. 2B), eight TMZ-resistant clones (A to H clones) were isolated. These clones were initially subjected to MTT viability tests, in which each clone was treated with high doses of TMZ (250 μM and 500 μM), to verify the increase in its resistance to chemotherapy (Fig. 2C). In conjunction with viability, live cell counts (cell growth curves) verified the proliferative profile of these cells over five consecutive days (Fig. 2D). These assays allowed the identification of three distinct resistant groups: the first group (lower cumulative doses and consistent treatments with small concentrations of TMZ), including clones A and B, maintained their proliferative capacity, but presented a very small increase in cell viability, and consequently, a phenotype of low resistance to TMZ; the second group (intermediate cumulative doses with crescent concentrations of TMZ), including clones C, D and E, presented an intermediate increase in viability, indicating a high resistance phenotype, besides to maintain the proliferative characteristic similar to the parental lineage; and the third group (higher cumulative doses and consistent treatments with high concentrations of TMZ), including clones F, G and H, presented a very high increase in resistance, but had a very high drop in the proliferative rate. Immunofluorescence assays of TMZ-resistant clones were performed to visualize the F-actin structure and cellular morphology, in addition to p53-Ser15 phosphorylation and p21^Kip1^ expression (Fig. 2E, 2F and S3). Morphological alterations in the F-actin structure appear to be dependent on the final cumulative dose, with distinct structures being observed and associated with viability assays (Fig. 2E). The first group (A and B) showed formation of stress fibers, but without very evident morphological changes. The actin structures observed in these cells are quite common in the stress response, indicating that the treatments did not promote abnormal changes in the activity of the Rho/actin pathway. The second group (C, D and E) presented a major change in the dynamics of F-actin, with formation of stress fibers, F-actin clusters and adhesion foci, and regions with high depolymerization. In addition, cell morphology was affected, with cells appearing more elongated, with formation of filopodia and protrusions indicating an unusually increased signaling of the Rho/actin pathway. The clones exposed to higher doses of TMZ (F, G and H) presented very drastic morphological changes, with many protrusions and cell fragments, polynucleated cells and very compromised F-actin dynamics, indicating again that, despite the high acquired resistance to TMZ, this group presents signs of extensive cell death. The expression of p53 and its phosphorylation at Ser15 were also affected in a manner dependent on the cumulative dose of TMZ (Fig. 2F). Clones B, C, D, E and F showed higher expression of p53 when compared with clones at the two extremes – lowest and highest cumulative dose – A, G and H, indicating that very attenuated or very extreme protocols for producing resistant cells either do not significantly activate or alter the p53 pathway, and both responses are not interesting for studying the mechanism here analyzed. Regarding p53 phosphorylation, clones exposed to more aggressive regimens showed higher levels of p-p53, which increased proportionally to the cumulative dose (Fig. 2F) and that perfectly reflected in the expression levels of p21^Kip^ (Fig. S3). This corroborates the phosphorylation data and the decrease in the proliferation rate presented by cells subjected to the most intense treatment cycles. These data indicate an extensive alteration of the p53 pathway, to a certain extent dependent on the final cumulative dose and the aggressiveness of the protocols adopted. Altogether, these analyses allowed us to exclude the clones obtained through the very attenuated protocols (A and B) and the most aggressive protocols (F, G and H), and only two remaining resistant clones, now called TMZ-R1 (formerly E) and TMZ-R2 (formerly D) were selected for subsequent characterization and experimentation.

**Figure 2.**
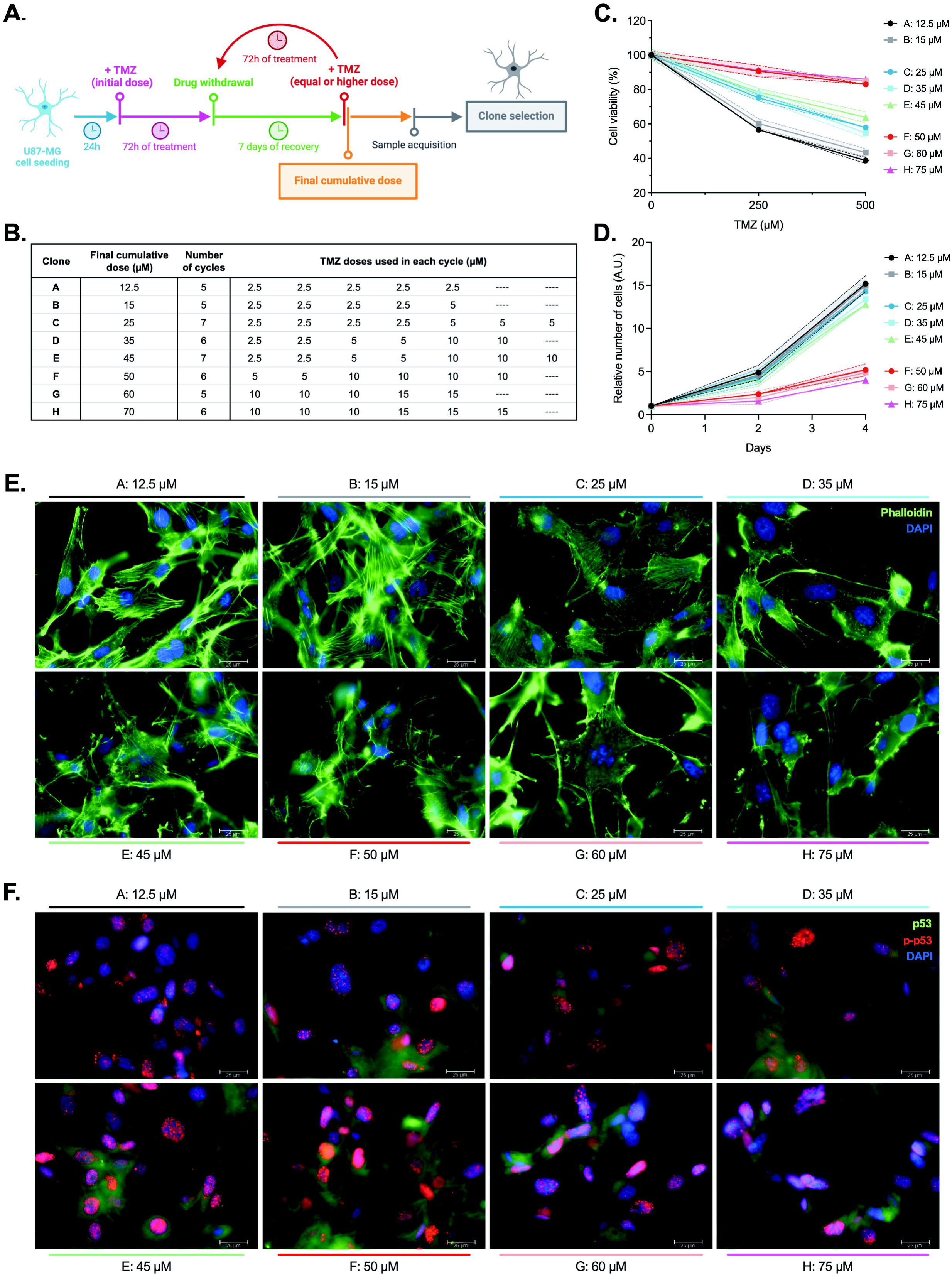
Proliferative capacity, F-actin structures, and the p53 pathway are differentially modulated by the TMZ treatment regimen during the acquisition of chemoresistance. **A.** Schematic representation of the protocol for generating TMZ-resistant U87-MG clones involving varying number of treatment/recovery cycles and final cumulative doses of TMZ. **B.** Final cumulative doses (µM), number of cycles, and TMZ doses used in each cycle for generating TMZ-resistant U87-MG clones (A–H). **C.** MTT viability assays of TMZ-resistant U87-MG clones (A–H) after TMZ treatment for 72 h. **D.** Cell growth curves of TMZ-resistant U87-MG clones (A– H). **E.** F-actin staining (green) of TMZ-resistant U87-MG clones (A–H). **F.** Immunofluorescence of TMZ-resistant U87-MG clones (A–H), showing p-p53 (Ser15) (red) and p53 (green). The images were captured with a 63x magnification objective and a scale bar of 25 μm.

### Characterization of TMZ-resistant U87-MG cell clones

Viability assays were performed to determine the IC_50_ for TMZ in the chosen chemoresistant clones to further determine the degree of resistance (Fig. 3A). Both clones showed a significant increase in resistance to TMZ: TMZ-R1 cells showed the highest degree of resistance to TMZ (IC_50_ of 929 μM), approximately 2 times of that obtained for the parental cells, while TMZ-R2 cells presented an intermediate resistance to TMZ (IC_50_ of 659 μM). Analysis of actin filament structure and cell morphology (Fig. 3B) showed that the TMZ-resistant clones present altered cell morphology, with increased formation of stress fibers, F-actin clusters, filopodia and lamellipodia, in addition to the formation of irregular protrusions (yellow arrows). Proliferation curves of the resistant clones treated with different doses of TMZ were compared to parental U87-MG cells under the same conditions (Fig. 3C). TMZ resistant clones showed a proliferation rate like U87-MG cells under non-stress conditions, with a doubling time of 1.9 days, which remain almost unaltered when exposed to 10 μM TMZ. Treatment with 25 μM or 50 μM TMZ reduced the cell proliferation and increased the doubling time of the parental cells of 2.3 and 3.8 days, respectively, while in resistant clones the loss of proliferation was smaller with doubling time of 2.1 and 3.0 days. Treatment with 50 μM TMZ had the strongest effect on the proliferation of cells: in parental U87-MG cells the proliferative capacity was almost eliminated, whereas in the resistant clones proliferation was reduced by 75% and did not prevent the cells from growing again in the last days. To corroborate morphological changes on actin cytoskeleton (Fig. 3B), pulldown assays to measure Rho activity were performed and showed that both resistant clones present increased levels of active RhoA, profilin 1 expression and in the phosphorylation of cofilin 1 compared to U87-MG cells (Fig. 3D). These data reinforce the Rho/actin pathway as one of the sensitive targets for the acquisition of TMZ resistance, just like what is observed for the acquisition of radioresistance (9). Another interesting feature observed for the TMZ-resistant clones, was the expression of different proteins involved in the repair of DNA damage specifically caused by TMZ (Fig. 3E). Both resistant clones have lower expression of MLH1 and MSH2 and a higher expression of ku80 and Rad51, indicating a decrease in the activity of the MMR repair pathway and an elevated strand break repair by NHEJ and HR, respectively. Altogether, these modulations can contribute to a lower sensitivity to TMZ (13). The modulation of the p53 pathway was also analyzed in the TMZ-resistant clones, regarding its expression, phosphorylation (Ser15), nuclear localization, and the expression of p21^Kip1^ (Fig. 3F-3I). In both resistant clones, the expression of p21^Kip1^ was decreased when compared to the parental U87-MG cells, with a greater effect in the resistant clone TMZ-R1 (Fig. 3F and 3H). Conversely, the expression of p53, as well as its phosphorylation, increased significantly in resistant clones (Fig. 3F and 3H), which showed lower localization of p53 in the nucleus (Fig. 3I). Last, but not least, the expression of MGMT was analyzed for these clones, a characteristic methyltransferase that promotes the direct repair of O^6^-MeG lesions caused by TMZ (Fig. 3G). Immunofluorescence analysis showed an increase in the expression of MGMT, quite diffuse in the nucleus of the TMZ-R clones (Fig. S4), subsequently confirmed by immunoblotting, showing that MGMT is one of the mechanisms positively regulated along with the acquisition of resistance to TMZ in our model.

**Figure 3.**
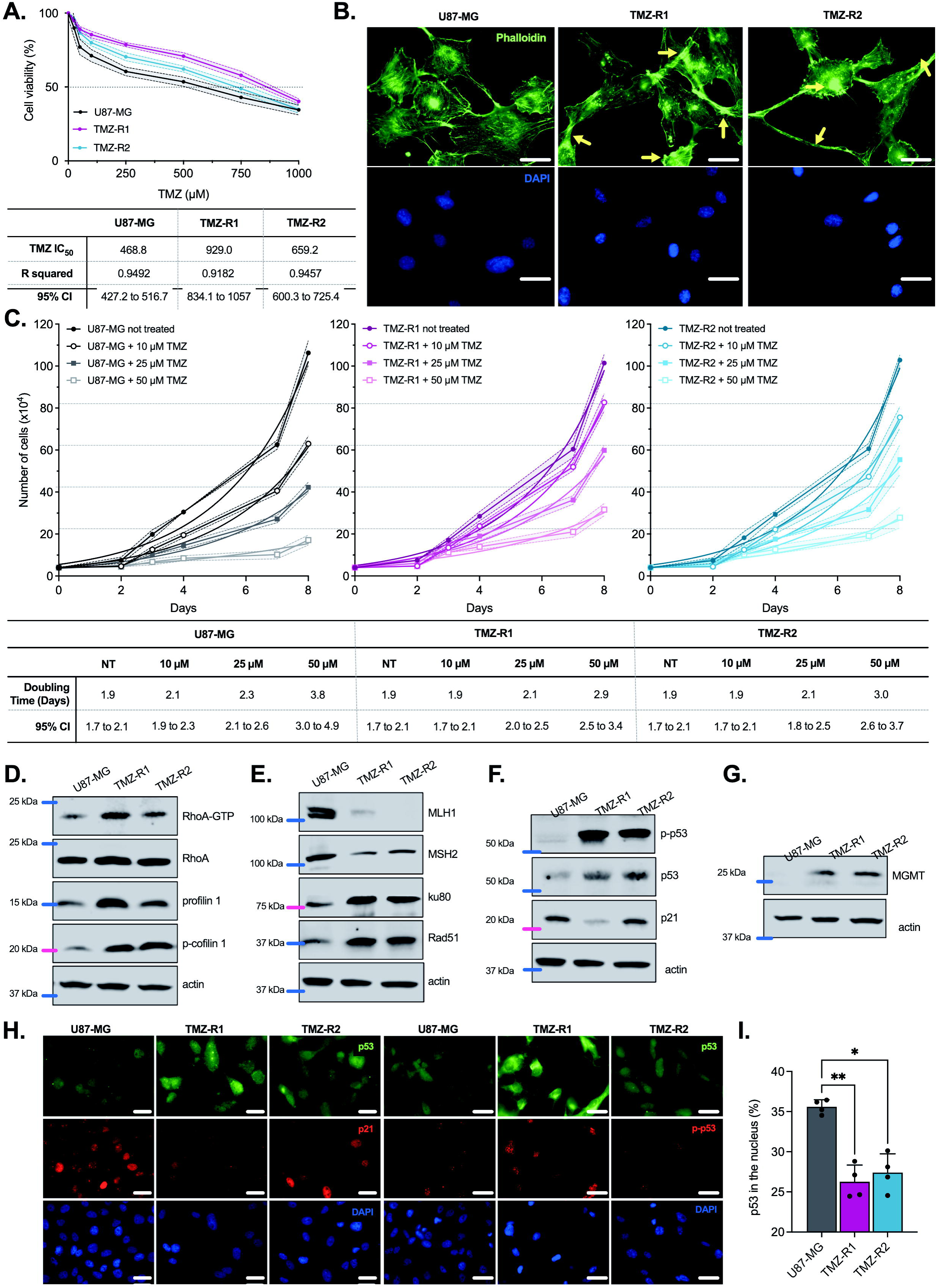
TMZ-resistant U87-MG clones display strong alterations on Rho/actin, DNA repair, and p53 pathways. **A.** MTT viability assays of U87-MG, TMZ-R1, and TMZ-R2 cells after TMZ treatment for 72 h. **B.** F-actin staining (green) of U87-MG, TMZ-R1, and TMZ-R2 cells. **C.** Cell growth curves of U87-MG, TMZ-R1, and TMZ-R2 cells following TMZ treatment. **D.** Representative image of pulldown assay for Rho activity measurement and immunoblotting showing the expression of profilin 1 and p-cofilin 1 on U87-MG, TMZ-R1, and TMZ-R2 cells. **E.** Immunoblotting of the indicated DNA repair-related proteins on U87-MG, TMZ-R1, and TMZ-R2 cells. **F.** Immunoblotting of p53 pathway proteins and MGMT on U87-MG, TMZ-R1, and TMZ-R2 cells. **G.** Immunofluorescence of U87-MG, TMZ-R1, and TMZ-R2 cells, showing p53 (green), p21 (red, left panel), and p-p53 (red, right panel). The images were captured with a 63x magnification objective and a scale bar of 25 μm. **H.** Nuclear p53 quantification of U87-MG, TMZ-R1, and TMZ-R2 cells.

### Rho inhibition abridged survival of TMZ-resistant cells

To verify whether inhibition of the Rho pathway could reverse the resistance phenotype acquired by TMZ resistant clones, we used the TMZ-R1 clone due to the highest IC_50_ for TMZ. Cell viability assays, under Rho inhibition by pEF-C3 transfection and further exposure to TMZ, reduced the IC_50_ of the TMZ-R1 cells by half, approaching the IC_50_ presented by three different wild-type p53 glioblastoma cell lines, U87-MG, U343-MG and A172 (Fig. 4A). Confirming that this mechanism works similarly to the previously one described for radiotherapy treatments (9), Rho inhibition did not modulate IC_50_ for TMZ (Fig. S5A) in p53-mutated GBM cells (T98G, U138-MG and U251-MG). By repeating these experiments treating cells with CP, Rho inhibition was able to sensitize parental and TMZ-R1 cells, similarly, reducing the IC_50_ for CP by more than half (Fig. 4B), nevertheless not affecting the IC_50_ for CP in p53-mutated GBM cells (Fig S5B). Corroborating cell viability data, clonogenic survival assays for the same set of cell lines under Rho pathway inhibition resulted in similar reduction of cell survival after exposure to TMZ (Fig. 4C) or CP treatments (Fig. 4D), with no evident effect in p53 mutated cells (Fig. S5C and S5D). To assess proliferation capacity of the TMZ-R1 clone in comparison to parental U87-MG cells, growth curves were performed combining Rho inhibition by C3 with TMZ or CP treatments. Doubling times were measured for U87-MG cells treated with 10 µM TMZ or 1.5 µM CP, while TMZ-R1 cells were treated with 25 µM TMZ or 2.5 µM CP (Fig. 4E and 4F). It is worth noting that Rho inhibition was an even better attenuator of TMZ-R1 clone proliferation compared to isolated drug treatments. Notably, the combination of C3 with either chemotherapy drugs resulted in higher doubling times and further reduced proliferative capacity of TMZ resistant cells.

**Figure 4.**
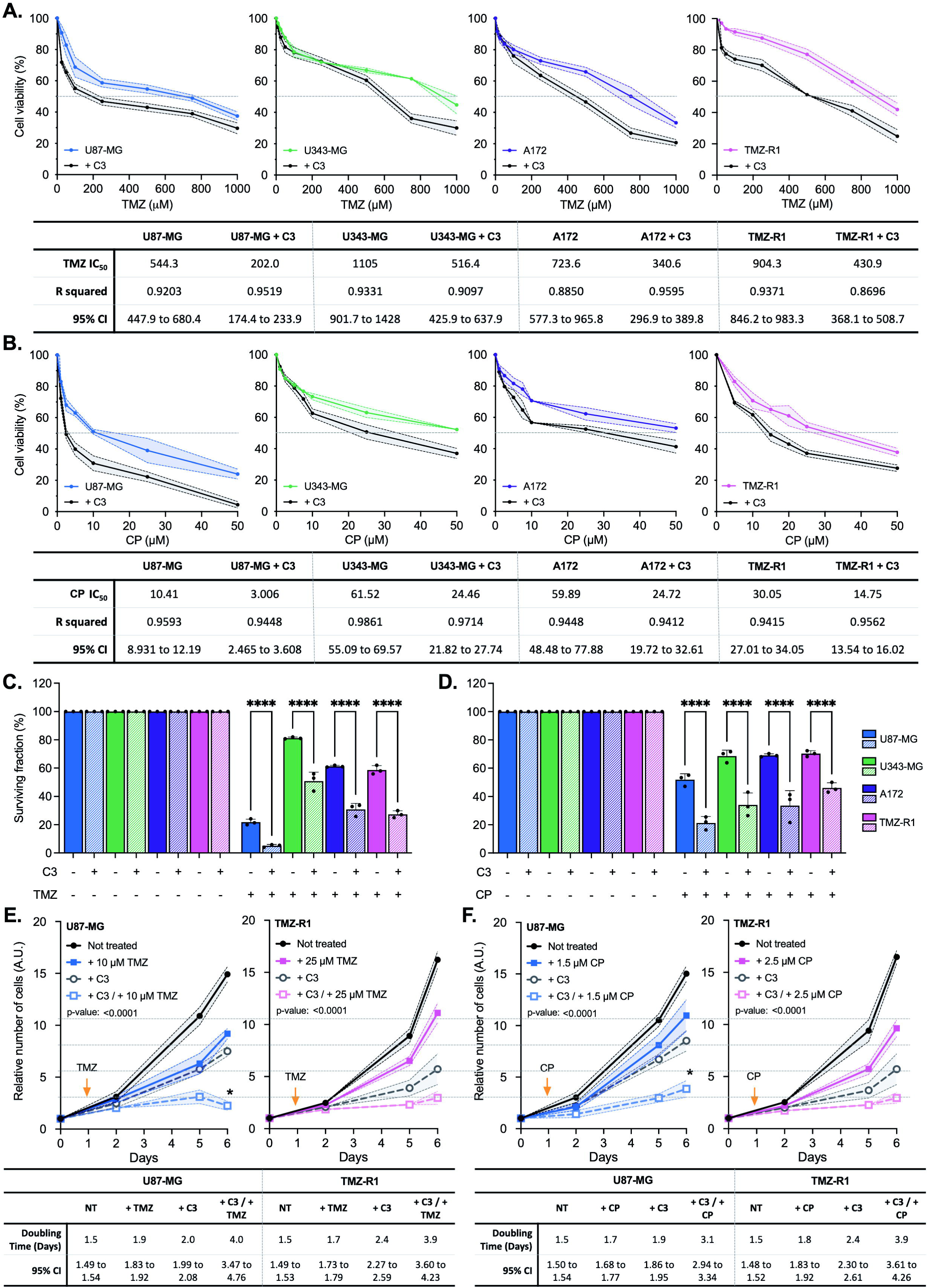
C3-mediated inhibition of Rho GTPase sensitizes wild-type p53 GBM cells to TMZ and CP treatments. **A.** MTT viability assays of wild-type p53 GBM cells – U87-MG, U343-MG, A172, and TMZ-R1 – transfected with C3 and treated with varying doses of TMZ for 72 h. **B.** MTT viability assays of wild-type p53 GBM cells transfected with C3 and treated with varying doses of CP for 72 h. **C.** Colony formation assays of wild-type p53 GBM cells transfected with C3 and exposed to 100 μM TMZ for 24h. **D.** Colony formation assays of wild-type p53 GBM cells transfected with C3 and exposed to 25 μM CP for 24h. **E.** Cell growth curves of U87-MG and TMZ-R1 cells transfected with C3 and treated with the indicated doses of TMZ for 24h. **F.** Cell growth curves of U87-MG and TMZ-R1 cells transfected with C3 and treated with the indicated doses of CP for 24h.

### C3-mediated inhibition of Rho GTPase attenuates the repair of TMZ-induced damage in TMZ-R1 cells

Rho pathway inhibition sensitize cells to chemotherapy drugs and TMZ resistant cells have increased activity of DNA repair pathways therefore suggesting that the Rho would be modulating DNA repair mechanisms in TMZ resistant cells. To assess this point, host-cell reactivation (HCR) assays were performed using U87-MG and TMZ-R1 cells under Rho inhibition by C3 toxin. In this assay, a luciferase reporter plasmid (pShuttle-Luc) was treated or not with TMZ or CP to promote specific DNA damage and further transfected into desired cells previously exposed to Rho inhibition (Fig. 5A). Luminescence measurements of luciferase expression and activity are directly correlated to DNA repair capacity of the remaining viable cells. As expected, TMZ treatment caused greater reduction in DNA repair of three parental cells (U87-MG, U343-MG and A172) than in the TMZ-R1 clone. However, all cell lines had their DNA repair capacity reduced around 20% by the C3 toxin (Fig. 5B), and the combination of C3 plus TMZ treatment attenuated DNA repair of the TMZ-R1 clone as well as U87-MG cells treated with TMZ alone. Of note, sensitization of TMZ resistant cells by Rho inhibition is dependent on wild type p53 expression, since downregulation of p53 by siRNA in U87-MG cells abolishes the C3 toxin effect on the reduction of DNA repair capacity (Fig. 5C). On the other hand, in cells with p53 mutations, Rho inhibition did not modulate the repair of TMZ-induced DNA lesions (Fig. S6A), though the reestablishment of wild type phenotype of p53-mutated U138-MG cells with PRIMA-1 (PR-1) restores this mechanism in the p53-mutated scenario (Fig. 5D). The DNA repair of the TMZ-R1 clone and wild type p53 cells is also affected by CP treatments and, again, all cells present reduced DNA repair capacity under Rho pathway inhibition (Fig. 5E). In agreement with TMZ results, the effect of Rho inhibition in U87-MG cells treated with CP is also dependent on wild type p53 background (Fig. 5F-5G and Fig. S6B).

**Figure 5.**
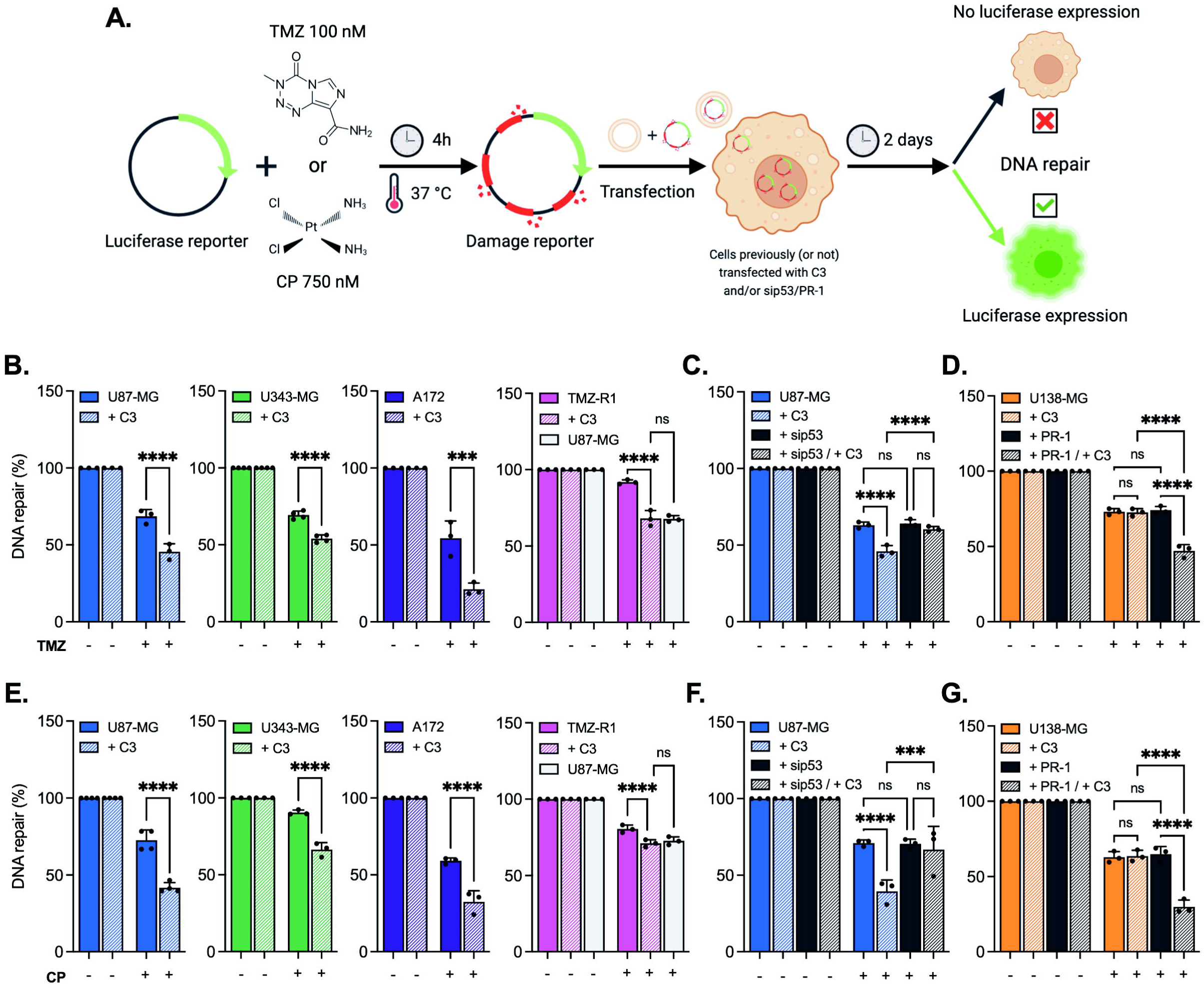
Inhibition of Rho GTPase attenuates the repair of TMZ- and CP-induced damage in TMZ-R1 cells dependent on the p53 background. **A.** Schematic representation of host-cell reactivation (HCR) assay. **B.** HCR assay of wild-type p53 GBM cells – U87-MG, U343-MG, A172, and TMZ-R1 – transfected with TMZ-treated luciferase reporter and subjected to C3-mediated Rho inhibition. **C.** HCR assay of U87-MG transfected with TMZ-treated luciferase reporter and subjected to C3-mediated Rho inhibition and p53 knockdown. **D.** HCR assay of U138-MG transfected with TMZ-treated luciferase reporter and subjected to C3-mediated Rho inhibition and p53 transcriptional activity restoration by PR-1. **E.** HCR assay of wild-type p53 GBM cells transfected with CP-treated luciferase reporter and subjected to C3-mediated Rho inhibition. **F.** HCR assay of U87-MG transfected with CP-treated luciferase reporter and subjected to C3-mediated Rho inhibition and p53 knockdown. **G.** HCR assay of U138-MG transfected with CP-treated luciferase reporter and subjected to C3-mediated Rho inhibition and p53 transcriptional activity restoration by PR-1.

### Rho pathway inhibition by disruption of actin cytoskeleton sensitizes chemoresistant spheroids

Looking for an approximation to the clinical scenario, an approach was developed for in vitro formation of tumor spheroids using U87-MG and TMZ-R1 cells. These spheroids were characterized as to the viability using acid phosphatase assay (APH) and further submitted to Rho pathway inhibition combined with chemotherapy drug treatments (Fig. 6A). Spheroid characterization involved: i) different numbers of seeding cells, and ii) time needed for their formation, being subsequently evaluated by morphology, diameter, sphericity and viability (Fig. 6B, 6C and 6D). Results show that the number of seeding cells, but not time of formation, reflects in the increased spheroid diameter for both cells, whereas their morphologies are very similar. These data are quite different from what we observed in spheroids constituted of radioresistant GBM cells (9). However, for parental and chemoresistant cells the viability is slightly different: parental U87-MG cells present increasing viability with time of formation despite the shrinking diameter; the TMZ-R1 clone remains viable along the days of formation even with diminishing diameter. Next, the Rho pathway inhibition was achieved by disrupting the actin cytoskeleton using Cytochalasin D (CD), a compound that inhibits actin polymerization or F-actin formation. By combining CD with TMZ treatment, cell viability of spheroids formed by U87-MG and TMZ-R1 cells were analyzed (Fig. 6E and 6F). The TMZ-R1 clone presented resistance to increasing doses of TMZ with very high IC_50_ in comparison to parental cells, nevertheless CD treatment reverted this resistant phenotype bringing down the IC_50_ for TMZ of this clone close to that of U87-MG cells, which was strongly sensitized by CD. When spheroids of these two cells were submitted to combination of CD with CP treatments, similar results were obtained (Fig. 6G and 6H) being TMZ-R1 even more sensitized to CP than TMZ, and with higher IC_50_ value. These results confirm that abrogation of Rho pathway and actin cytoskeleton can reverse chemoresistance of glioblastoma cells for TMZ and CP drugs.

**Figure 6.**
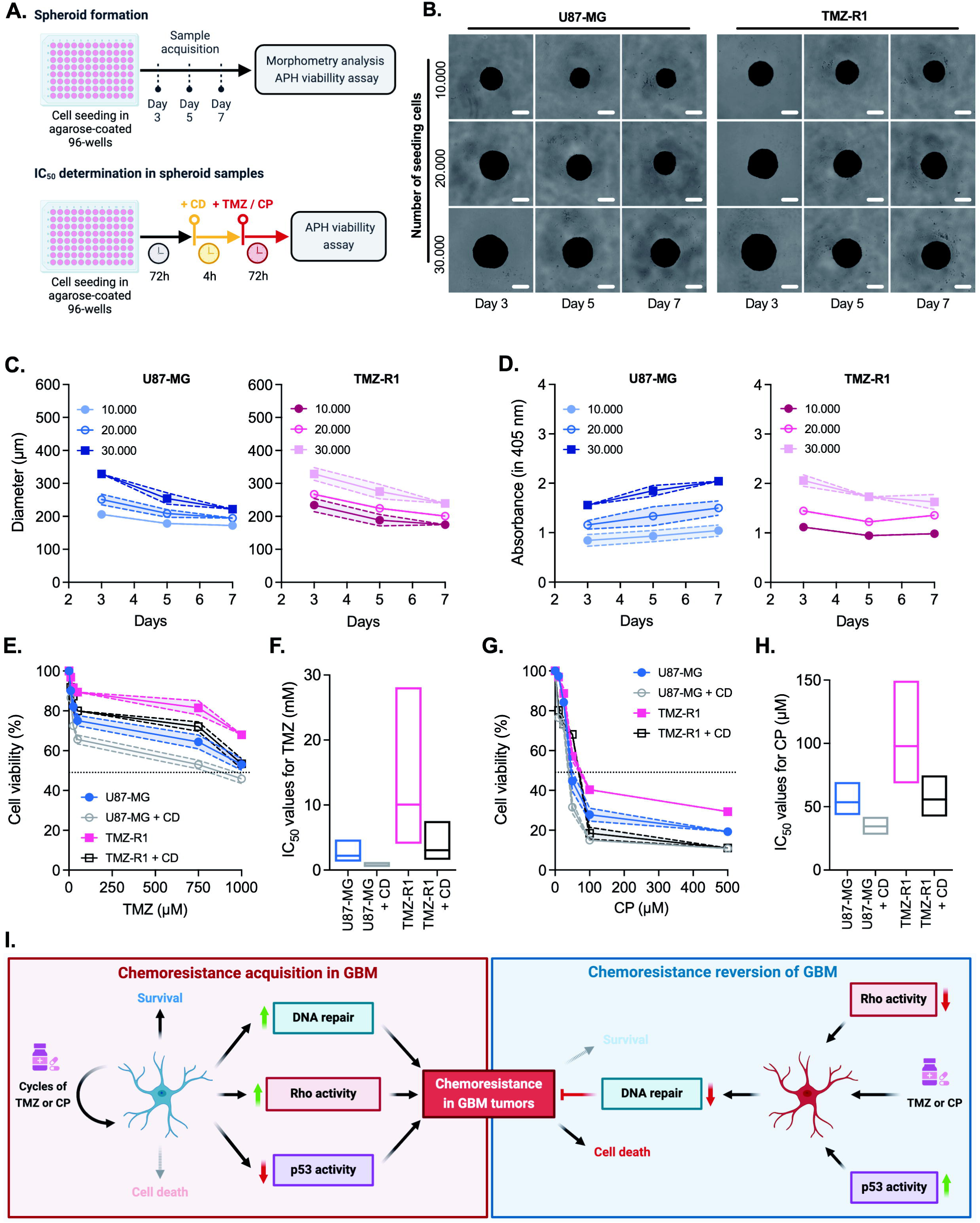
Pharmacological inhibition of Rho pathway sensitizes chemoresistant spheroids to TMZ and CP treatments. **A.** Schematic representation for experiments using U87-MG- and TMZ-R1-derived spheroids, indicating the time-point for treatment and/or data acquisition for each specific assay. **B.** Micrographs of spheroids obtained from U87-MG and TMZ-R1 with varying initial numbers of cells and formation time. The images were captured with a 5x magnification objective and a scale bar of 100 μm. **C.** Diameter quantification of spheroids obtained from U87-MG and TMZ-R1 showed in (**B**). **D.** Cell viability assessment by APH assay of spheroids obtained from U87-MG and TMZ-R1 showed in (**B**). **E.** APH viability assays of spheroids obtained from U87-MG and TMZ-R1 cells pre-treated with CD 100 nM and treated with varying doses of TMZ for 72 h. **F.** IC_50_ values for TMZ of spheroids obtained from U87-MG and TMZ-R1 cells pre-treated with CD. **G.** APH viability assays of spheroids obtained from U87-MG and TMZ-R1 cells pre-treated with CD 100 nM and treated with varying doses of CP for 72 h. **H.** IC_50_ values for CP of spheroids obtained from U87-MG and TMZ-R1 cells pre-treated with CD. **I.** Schematic model of GBM chemoresistance acquisition and reversion by targeting of the Rho and p53 pathways.

## DISCUSSION

Cancer cells are usually growing and dividing in an uncontrolled manner compared to normal cells and thereby possess very high levels of endogenous stress. Chemotherapeutic treatments work by inhibiting the growth and division of cancer cells, rapidly and more effectively destroying tumor cells without affecting the non-cancerous cells (14). Over the years, numerous anti-cancer drugs and natural medicinal compounds have been developed to suppress tumor growth through various mechanisms, including the ability to interfere with critical cellular processes, such as programmed cell death/apoptosis, DNA replication, immune response, and DNA repair (3,15). In the GBM scenario, both TMZ and CP are highly effective in promoting cytotoxicity to these tumors. However, both chemotherapeutic agents are frequently subject to mechanisms of chemoresistance, making them suitable models for our chemoresistance study. TMZ-based chemotherapy is undoubtedly the most widely used and effective treatment currently applied in the clinic. Nevertheless, there is growing interest in developing new therapeutic regimens for GBM, where TMZ has been combined with various other cytostatic or cytotoxic agents to overcome or minimize chemoresistance. Over the past decade, numerous phase I and II studies have explored the safety and efficacy of TMZ combined with other novel or conventional chemotherapeutics, including CP, for the treatment of recurrent GBM tumors (16,17). The work here discussed sheds new light on the understanding of glioblastoma chemoresistance, revealing a mechanism of multiple resistance acquisition to CP and TMZ mediated by the Rho pathway. Moreover, the results suggest the potential use of this mechanism for sensitization of chemoresistant GBM to increase the efficacy of these treatments.

TMZ and CP are two very distinct molecules that promote genotoxic effects by generating different types of DNA lesions. TMZ is an imidazotetrazine lipophilic prodrug that is rapidly and nonenzymatically converted to the active alkylating metabolite MTIC [(methyl-triazene-1-yl)-imidazole-4-carboxamide], which cytotoxic effects are DNA methylation and production of the O^6^-methylguanine (O^6^-MeG) lesions that lead to mismatches during replication, ultimately resulting in DSBs (16). CP, on the other hand, exerts its cytotoxic effect through the formation of DNA adducts on double-stranded DNA, leading to cell cycle arrest and eventual cell death (18). The effectiveness of these chemotherapeutic agents is counteracted by robust DNA repair mechanisms. TMZ predominantly triggers the mismatch repair (MMR) pathway, while CP activates the nucleotide excision repair (NER) pathway.

Our group recently elucidated a mechanism showing that Rho/actin modulates GBM sensitivity to ionizing radiation by regulating DSB repair mechanisms through the interaction and nuclear translocation of p53 mediated by G-actin, which also affects DDR signaling (9). This mechanism can also be applied in this work, since our data showed that Rho inhibition drastically impairs the ability of GBM cells to repair TMZ-induced lesions and CP adducts only in a wild-type p53-dependent manner. Indeed, TMZ preferentially induces cell cycle arrest at the G2/M phase, in a mechanism that appears to be independent of p53 status but has a much more pronounced effect in GBM cells expressing wild type p53 (19). It is also known that the Syx-RhoA-Dia1-YAP/TAZ signaling axis regulates cell cycle progression and DNA damage repair in response to TMZ in primary GBM culture models (20), while TMZ treatment of U251-MG cells (mutant p53 status) did not affect Rho GTPase activity (21). Regarding CP adducts repair, the NER pathway is the most well-known mechanism for removing this type of damage and has also been shown to be regulated by RhoA activity under UV-induced DNA damage (22). Similarly, in this work, we observed that Rho inhibition affected the response to TMZ with a much greater impact on the wild type p53 cell line. This observed effect may be attributed to the suppression of DDR promoted by Rho inhibition, since DNA lesions induced by TMZ promote replication forks to stall, initially activating the ATR-Chk1 axis of the DNA damage response (DDR) pathway, followed by the activation of the ATM-Chk2 axis due to the secondary formation of DSBs at collapsed replication forks (13,16).

Chemotherapy drugs are usually administered at regular, repeated intervals known as treatment cycles. The timing of these cycles is based on the ability of normal tissues to recover and is kept as short as possible. Although some repopulation occurs between cycles, tumors have a reduced capacity of repair compared to normal tissues and, as a result, repeated chemotherapy cycles gradually reduce the tumor population and allow normal cells to recover during the intervals (14,16). In GBM tumors, TMZ offers superior benefits since it has a higher ability to cross the blood-brain barrier compared to other cytotoxic drugs, and a higher patient tolerance, since gliomas present a more alkaline environment compared to surrounding healthy brain cells, leading to drug activation almost exclusively in the tumor cells (1,23). Nevertheless, repeated chemotherapy cycles often lead to chemoresistance, as shown in our results, where just two cycles of low-dose TMZ or CP significantly increased GBM cell tolerance. Surprisingly, we observed that the treatments cycles increase not only the expression and activity of DNA repair proteins such as MGMT, p53, Rad51, and Ku80 (and decrease the activity of mismatch repair proteins, as expected), but also the activity of Rho pathway as well as actin polymerization. Moreover, differential gene expression analyses in U87-MG cells treated with low doses of CP over different exposure times showed that short CP exposure modulates p53 signaling and DNA-repair pathways, whereas long exposure modulates not only p53 pathways but also Rho GTPase pathway, indicating signaling from p53 to Rho GTPases. These data strongly suggest that the process involved in the acquisition of chemoresistance involves the positive modulation of Rho/actin pathways together with DNA repair pathways.

The acquisition of chemoresistance in gliomas is multifactorial and involves, beyond the increase in DNA repair activity, other mechanisms such as increased production of reactive oxygen species (ROS) (24), the presence of glioma stem cells (GSCs) (25,26), dysregulated microRNAs (27,28), altered metabolism (29), c-Met signaling (30), and ATP-binding cassette (ABC) transporters (31). These factors enhance tumor survival under chemotherapy by promoting DNA repair, drug efflux, and other protective mechanisms, facilitating resistance to DNA-damaging agents (16). The role of the Rho pathway in the acquisition and maintenance of chemoresistance in different types of tumors have emerged in the past decade, such as in breast (32), ovary (33), cervical (34), lung (35), liver (36,37), pancreatic (38), among others. It was recently shown that cisplatin-resistant human squamous carcinoma cells exhibit higher expression of β-actin and more defined F-actin structures. Additionally, this study demonstrated that inhibition of actin cytoskeleton polymerization via CD increases the sensitization of these cells by altering chloride channels (39). It was also shown that the Rho GAP ARHGAP21 (which promotes Rho inactivation), considered a potent tumor suppressor, is more highly expressed in gliomas with a hypermethylated MGMT promoter, indicating that the expression of this GAP is associated with lower MGMT expression and increased sensitivity to chemotherapeutic agents, particularly TMZ (40). Additionally, our group has observed that inhibiting the Rho GTPase pathway can restore sensitivity to genotoxic therapeutic agents like ionizing radiation (IR), TMZ and CP, particularly in GBM cells with wild type p53 status, underscoring this pathway as a potential therapeutic target (41).

Another particularly interesting point of the study was the multiple resistance exhibited by cells continuously exposed to TMZ treatments, which also developed resistance to CP. In vitro studies have shown that CP can reduce MGMT activity, promoting increased resistance to TMZ (17), similar to the observations in the presented data, where continuous exposure to TMZ led to greater resistance to high doses of CP. All these data were further corroborated in GBM spheroid models, in which the pharmacological disruption of F-actin by CD also increased the sensitivity of both U87-MG and TMZ-resistant cells to TMZ and CP, reinforcing the mechanism here verified.

In summary, our data suggest that cycles of TMZ or CP treatments promotes the activation of Rho/actin pathway together with DNA repair pathways activation and p53 signaling inhibition (in a p53 wild type scenario). These modulations collectively induce the acquisition of GBM chemoresistance (Fig. 6I). When chemoresistant GBM cells are treated with TMZ or CP under p53 activation, Rho pathway inhibition leads to sensitization to chemotherapeutic drugs by downregulating DNA repair capacity, thus inhibiting chemoresistance in GBM (Fig. 6I). Therefore, this mechanism emerges as a good and promising candidate for targeting resistant GBM tumors in clinical approaches.

## DECLARATIONS

### Data availability statement

The databases used in bioinformatic analysis are available in online platforms, as describe in Material and Methods section. All data are available upon request from the corresponding author.

## Authors Contribution

Conception and design: Y.T. Magalhaes, F.L. Forti

Development of methodology: Y.T. Magalhaes, F.L. Forti

Acquisition of data: Y.T. Magalhaes

Analysis and interpretation of data: Y.T. Magalhaes, F.L. Forti, V.K. Boell

Writing, review, and/or revision of the manuscript: Y.T. Magalhaes, F.L. Forti, V.K. Boell

Administrative, technical, or material support: Y.T. Magalhaes, F.L. Forti

Study supervision: F.L. Forti

## Acknowledgments

The authors thank Prof. Elza T. S. Hojo, from the Pharmaceutical Science Faculty of Ribeirão Preto – University of São Paulo, as well as Prof. Carlos F. M. Menck and Dr. Veridiana Munford, from the Institute of Biomedical Sciences – University of São Paulo, for the donation and validation of the cell lines and for the support with the cytometer analysis. We thank Prof. Alexandre B. Cardoso and Prof. Nicolas C. Hoch, from the Institute of Chemistry – University of São Paulo, for the use of microscopes Leica DMi8 widefield and Zeiss LSM 780 confocal, respectively. We also thank Prof. Bianca S. Zingales and the technician Marcelo N. Silva, from the Institute of Chemistry - University of São Paulo, for equipment usage and reagents exchange.

