## Supplementary Tables for "Acquisition and reversal of glioblastoma chemoresistance are mediated by the Rho GTPase pathway"

### Supplementary Data

**Supplementary Table S1.** Antibodies incubation conditions for immunobloting assay

| Antibody | Dilution | Incubation | Source | Company | CAT number |
|---|---|---|---|---|---|
| **Actin** | 1:2000 | 2 h, RT | *Goat* | Santa Cruz Biotechnology | sc-10731 |
| **phospho-cofilin 1 (Ser3)** | 1:500 | *Overnight*, 4°C | *Mouse* | Santa Cruz Biotechnology | sc-271921 |
| **Ku80** | 1:500 | *Overnight*, 4°C | *Goat* | Santa Cruz Biotechnology | sc-1485 |
| **MGMT** | 1:1000 | *Overnight*, 4°C | *Rabbit* | Cell Signalling Technology | #58121S |
| **MLH1** | 1:500 | *Overnight*, 4°C | *Rabbit* | Santa Cruz Biotechnology | sc-582 |
| **MSH2** | 1:500 | *Overnight*, 4°C | *Rabbit* | Santa Cruz Biotechnology | sc-494 |
| **p21** | 1:1000 | *Overnight*, 4°C | *Rabbit* | Santa Cruz Biotechnology | sc-397 |
| **p53** | 1:1000 | *Overnight*, 4°C | *Mouse* | Santa Cruz Biotechnology | sc-56180 |
| **phospho-p53 (Ser15)** | 1:1000 | *Overnight*, 4°C | *Rabbit* | Cell Signalling Technology | #9286 |
| **profilin 1** | 1:1000 | *Overnight*, 4°C | *Mouse* | Santa Cruz Biotechnology | sc-137235 |
| **Rad51** | 1:100 | *Overnight*, 4°C | *Mouse* | Santa Cruz Biotechnology | sc-398587 |
| **RhoA** | 1:500 | 4 h, RT | *Mouse* | Santa Cruz Biotechnology | sc-418 |

**Supplementary Table S2.** Antibodies incubation conditions for immunofluorescence assays

| Antibody | Dilution | Incubation | Source | Company | CAT number |
|---|---|---|---|---|---|
| **Alexa Fluor 488** | 1:500 | 1 h, RT | --- | Thermo Fischer Scientific | --- |
| **Alexa Fluor 555** | 1:500 | 1 h, RT | --- | Thermo Fischer Scientific | --- |
| **Alexa Fluor 568** | 1:500 | 1 h, RT | --- | Thermo Fischer Scientific | --- |
| **Alexa Fluor 647** | 1:500 | 1 h, RT | --- | Thermo Fischer Scientific | --- |
| **p21** | 1:300 | 3 h, RT | *Rabbit* | Santa Cruz Biotechnology | sc-397 |
| **p53** | 1:300 | 3 h, RT | *Mouse* | Santa Cruz Biotechnology | sc-56180 |
| **phospho-p53 (Ser15)** | 1:300 | 3 h, RT | *Rabbit* | Cell Signalling Technology | #9286 |
| **Phalloidin Alexa Fluor 488** | 1:1000 | 2 h, RT | --- | Thermo Fischer Scientific | A12379 |
