## Supplementary Figures for "Acquisition and reversal of glioblastoma chemoresistance are mediated by the Rho GTPase pathway"

### FIGURE S1

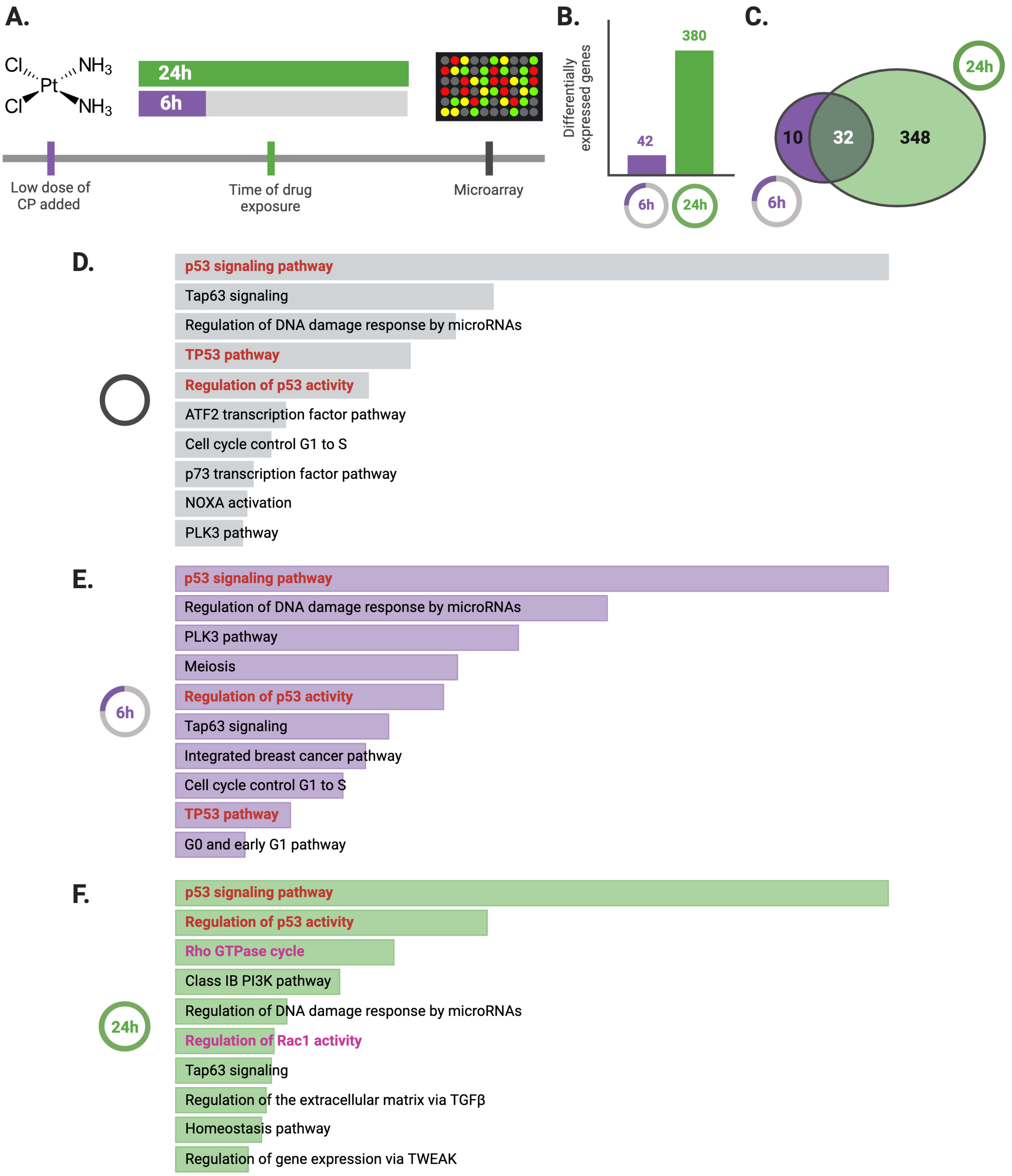
A.
Cl
Cl
Pt
NH3
NH3
24h
6h
Low dose of CP added
Time of drug exposure
Microarray
B.
Differentially expressed genes
380
42
6h
24h
C.
24h
10
32
348
6h
D.
p53 signaling pathway
Tap63 signaling
Regulation of DNA damage response by microRNAs
TP53 pathway
Regulation of p53 activity
ATF2 transcription factor pathway
Cell cycle control G1 to S
p73 transcription factor pathway
NOXA activation
PLK3 pathway
E.
6h
p53 signaling pathway
Regulation of DNA damage response by microRNAs
PLK3 pathway
Meiosis
Regulation of p53 activity
Tap63 signaling
Integrated breast cancer pathway
Cell cycle control G1 to S
TP53 pathway
G0 and early G1 pathway
F.
24h
p53 signaling pathway
Regulation of p53 activity
Rho GTPase cycle
Class IB PI3K pathway
Regulation of DNA damage response by microRNAs
Regulation of Rac1 activity
Tap63 signaling
Regulation of the extracellular matrix via TGFβ
Homeostasis pathway
Regulation of gene expression via TWEAK

**Supplementary Figure S1. CP exposure affects the expression of genes belonging to the Rho GTPase and p53 pathways in U87-MG GBM cells. A.** Bioinformatics analysis conducted using experimental data available online from the EMBL Cell Expression Atlas platform, where U87-MG cells were treated with CP for 6h and 24h, and their transcriptome was analyzed by Microarray. The raw expression data were obtained in log2(Fold change) between treated conditions and the baseline expression of the cell line. **B.** Number of differentially expressed genes of CP-treated U87-MG cells for 6h and 24h. **C.** Venn diagram showing the intersection of differentially expressed genes between the two CP exposure times. Data were filtered with a threshold of p-value ≤0.05 and |log2(Fold change)| ≥ 1.00. **D-F.** Pathway enrichment analysis performed using the EnrichR platform with the dataset of differentially expressed genes in the U87-MG cell line independently of CP exposure time (**D**) or after 6h (**E**) and 24h (**F**) of CP exposure, showing the top 10 modulated pathways. The graphs were ranked according to the p-value of enrichment, with bar sizes relative to these values. Pathways common to the three groups are highlighted in bold, while pathways related to Rho GTPases are highlighted in red.

### FIGURE S2

A.

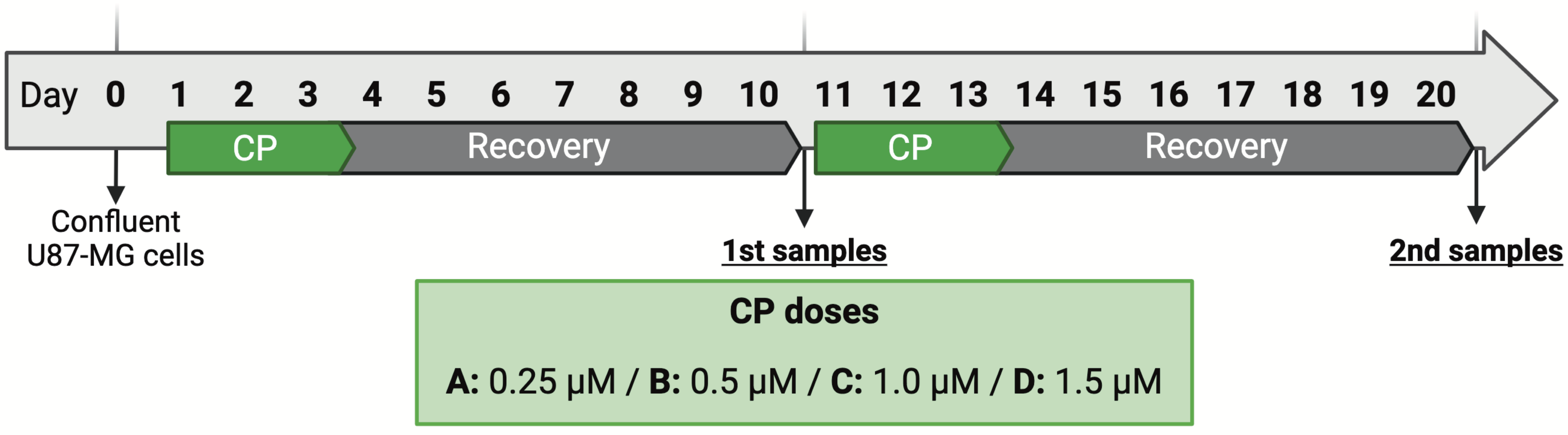


B.

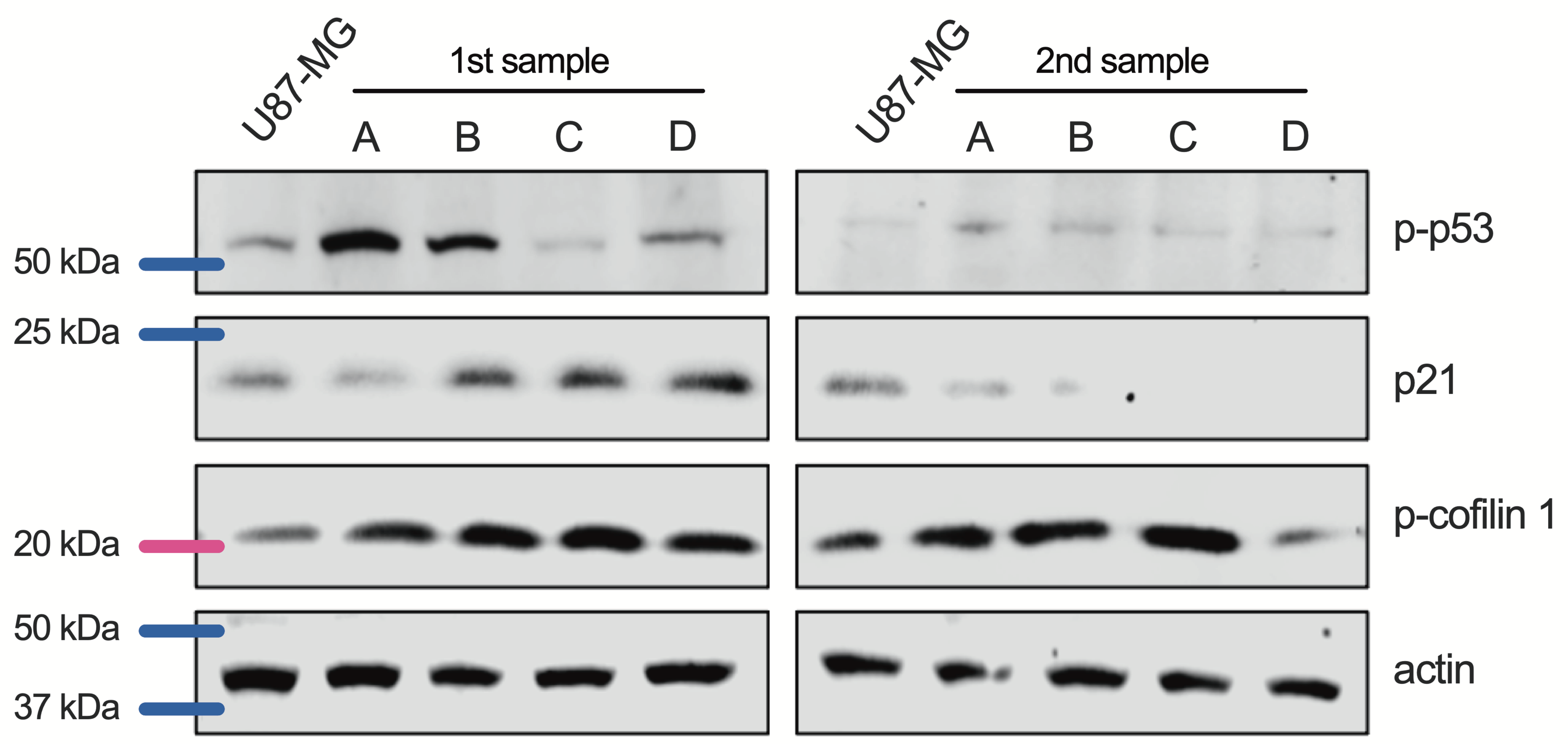


**Supplementary Figure S2. Rounds of CP exposure attenuate the p53 pathway and promote increased Rho pathway activity. A.** Schematic representation of treatment protocols involving two rounds of treatment/recovery with varying doses of TMZ in U87-MG cells for the specified duration. Cells were divided into groups (A–D) according to the indicated drug concentrations. Cells were harvested after the first (1st samples) and second rounds of treatment (2nd samples) and subjected to immunoblotting assays. **B.** Representative image of immunoblotting assays from U87-MG cell groups exposed to CP showing the expression of p21 and phosphorylation of p53 and cofilin 1 (Ser3).

### FIGURE S3

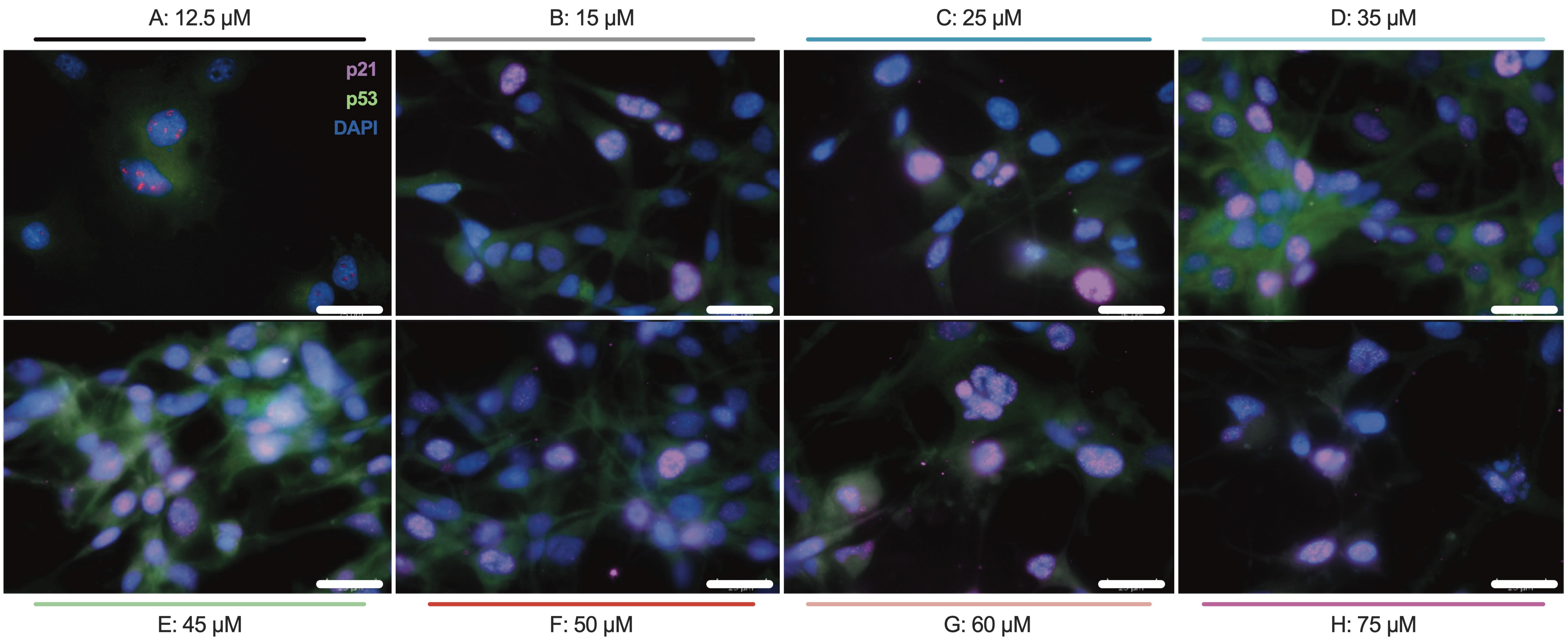


**Supplementary Figure S3. p53 signaling in TMZ-resistant U87-MG cells.** Immunofluorescence assays of each TMZ-resistant U87-MG cell clone obtained after the TMZ resistance acquisition protocol, showing p53 in green, p21 in pink, and DAPI in blue. The images were captured with a 63x magnification objective and a scale bar of 25 μm.

### FIGURE S4

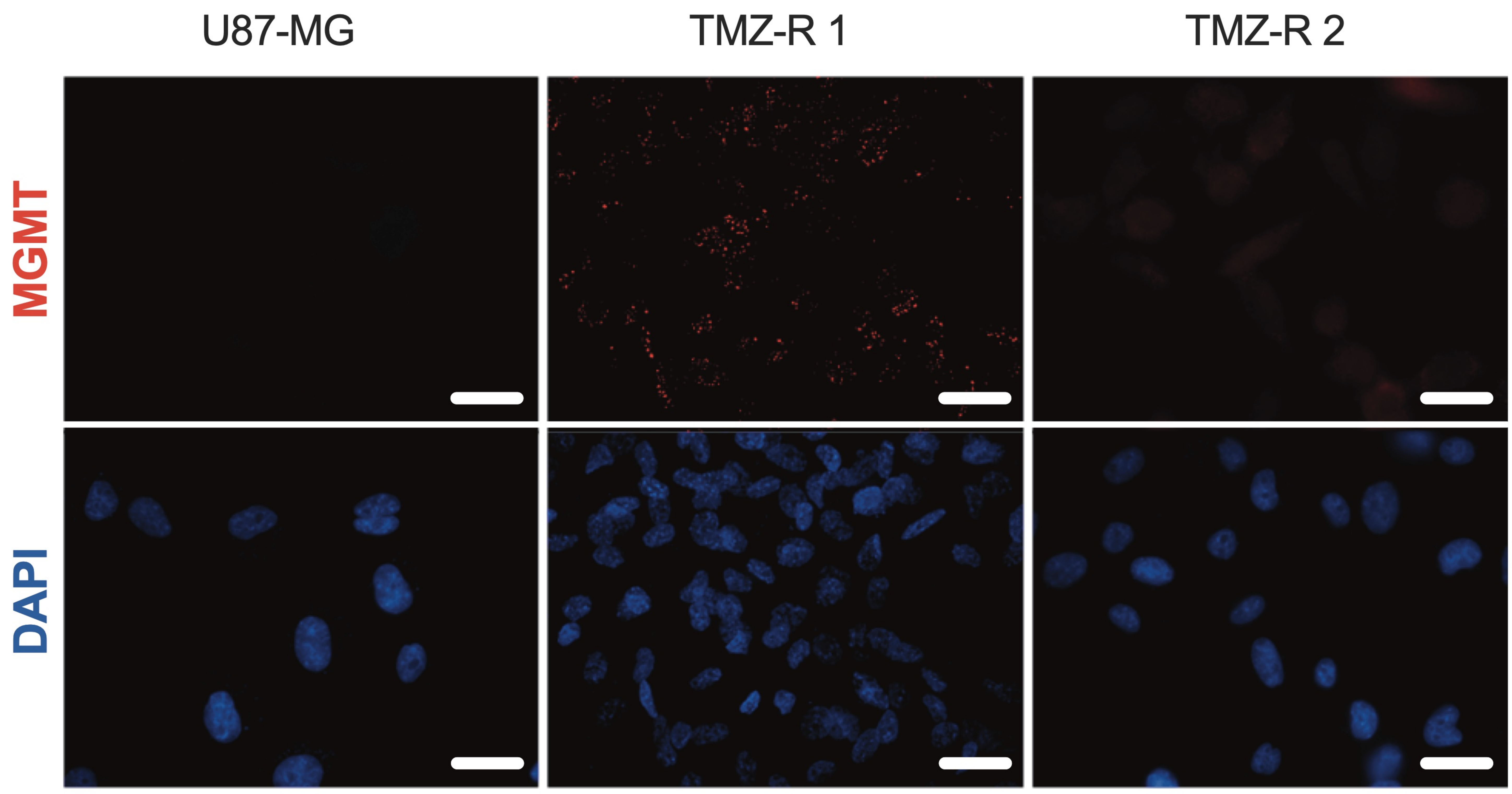


**Supplementary Figure S4. MGMT expression is increased in TMZ-resistant cells.** Immunofluorescence assay for characterization of TMZ-R1 and TMZ-R2 clones, showing MGMT in red and DAPI in blue. The images were captured with a 63x magnification objective and a scale bar of 25 μm.

### FIGURE S5

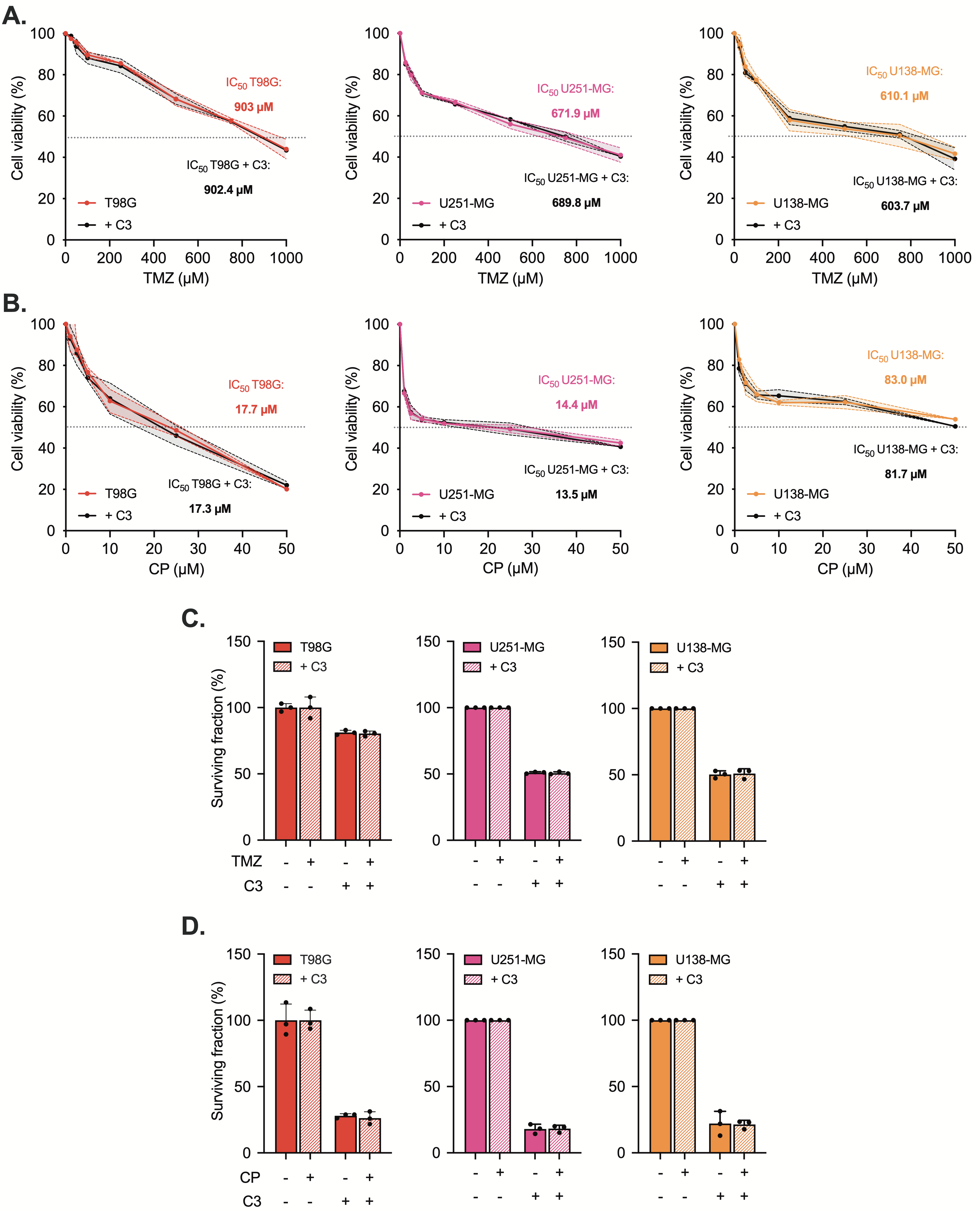


**Supplementary Figure S5. Rho inhibition does not affect the cellular response of mut-p53 GBM cells to TMZ or CP. A-B.** MTT viability assays of mut-p53 GBM cells—T98G, U251-MG, and U138-MG—transfected with C3 and exposed to different doses of TMZ (**A**) or CP (**B**). **C-D.** Colony formation assays of mut-p53 GBM cell lines exposed to 100 μM of TMZ (**C**) or 25 μM of CP (**D**).

**FIGURE S6**

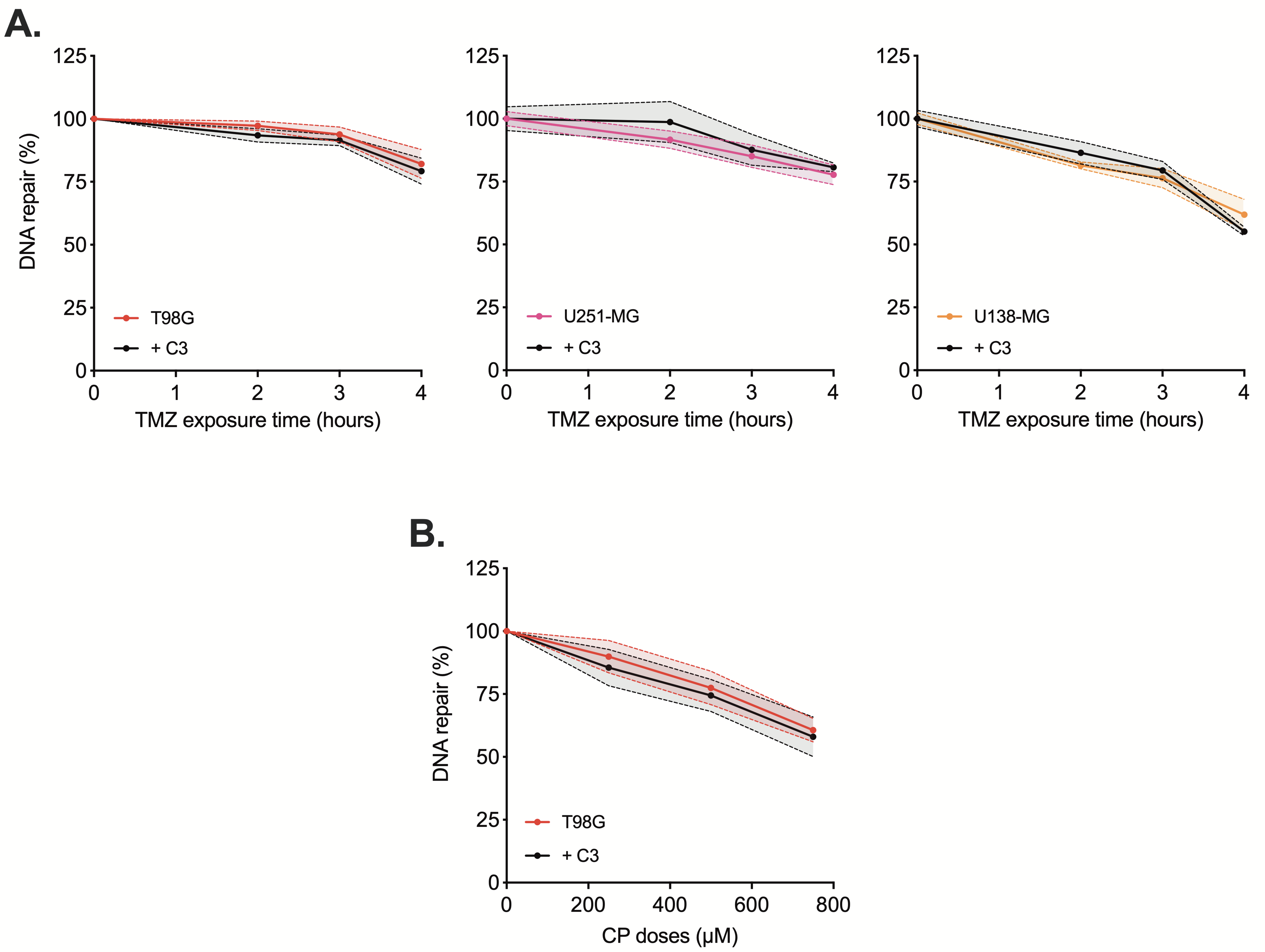


**Supplementary Figure S6. Rho inhibition does not affect the repair capacity of DNA lesions induced by TMZ or CP in mut-p53 GBM cells. A.** HCR assay of mut-p53 GBM cells – T98G, U138-MG and U251-MG – transfected with pShuttle-Luc plasmids treated with 100 nM TMZ at increasing incubation times, under C3-mediated Rho inhibition. **B.** HCR assay of mut-p53 GBM cells transfected with pShuttle-Luc plasmid with increasing doses of CP for 4h at 37ºC, and also subjected to Rho inhibition via C3.
